# Feasibility and safety of a telemetric pulmonary artery pressure monitoring system in acute and chronic porcine models of pulmonary hypertension

**DOI:** 10.1101/2020.12.27.424411

**Authors:** Alexander M. K. Rothman, Nadine D. Arnold, Jacob Abou-Hanna, Omid Forouzan, Andrew J. Swift, Payman Dahaghin, Shiran Konganige, Jennifer T. Middleton, Hamza Zafar, S. Kim Suvarna, David G Kiely, Julian Gunn

## Abstract

**Aims:** Pulmonary hypertension (PH) is associated with significant morbidity and mortality and leads to progressive right heart failure. In patients with PAH, haemodynamic parameters measured at catheterisation relate to clinical worsening events, in patients with heart failure proactive pulmonary artery pressure based therapeutic intervention reduces hospitalisation. We therefore investigated use of a novel implanted pulmonary artery (PA) pressure monitor to detect clinically relevant changes in pressure in large animal models of pulmonary hypertension (PH).

**Methods and Results:** Prototype pulmonary artery pressure sensors (Endotronix) were implanted using standard interventional techniques. Acute PH was induced by infusion of thromboxane A2 in domestic swine. Over a physiological range pressure monitors remained concordant to reference catheter (bias −0.43, 95%CI-5.3-4.4). Chronic PH was induced by i.p. injection of monocrotaline. Implanted pressure sensors demonstrated a gradual rise in PA pressure over 30 days (baseline: 20.7+/-0.4 vrs day-30: 31.74+/-1.4, p<0.01). Pressure sensor derived readings matched reference catheter at baseline and day-30. Pressure sensors remained stable and no adverse events were identified by clinical and histological examination.

**Conclusions:** The development of PA pressure monitors provide long-term haemodynamic data that identified clinically meaningful changes in pulmonary artery pressure. In addition to proactive heart failure management, such devices may be used to optimise or personalize patient therapy, investigate aspects of physiology and pathology essential to the understanding of disease and provide the opportunity to assess therapeutic interventions in clinical studies.

## Introduction

Pulmonary hypertension (PH) comprises a range of diseases defined by a resting mean pulmonary artery pressure (PAP) of ≥25 mmHg.^1-3^ Approved therapies are licensed only for patients with pulmonary arterial hypertension (PAH) and chronic thromboembolic pulmonary hypertension (CTEPH) acting through pharmacological modulation of the prostacyclin, endothelin or nitric oxide pathways.^1-3^ Excepting patients with a vasodilator response, drug choice is empirical and clinical management uniform, based upon an algorithm that matches functional class to a number of vasodilator agents.^2^ In clinical practise PAH progression is determined by assessment of symptoms, exercise capacity and measurements of right ventricular function^2,4^. Clinical studies of therapies for PH use a range of surrogate measures as primary endpoints including invasive measurement of pulmonary haemodynamics, 6-minute walk test, quality of life measures, non-invasive imaging and, in recent phase III studies, composite endpoints of clinical worsening.^5–9^ In patients with PAH observational studies have identified relationships between haemodynamic parameters measured at right heart catheterisation and clinical worsening events. Pooled patient-level data from over 1100 patients enrolled into FDA drug approval studies demonstrated that active treatments alter mean right atrial pressure (RAP), mean pulmonary artery pressure (mPAP), cardiac output (CO), cardiac index (CI), pulmonary vascular resistance (PVR) and pulmonary artery compliance, and that with knowledge of treatment allocation these parameters are predictive of clinical events.^10^ However, the invasive nature and cost of cardiac catheterisation usually limits its use to establishing diagnosis, follow-up in selected patients and the conduct of phase II studies.

The development of implantable pressure monitors permits alteration of clinical therapy based on daily pulmonary artery pressure measurements. The benefit of such hemodynamic parameter-guided therapy has been studied in patients with heart failure with both preserved and reduced ejection fraction. In these patient groups, elevation of cardiac filling pressures precedes clinical worsening events by ≥ 1 to 2 weeks.^11,12^ Early, proactive pressure-guided heart failure management using implantable pulmonary artery pressure monitors has been demonstrated to reduce hospitalisation in comparison to standard care.^13–15^ These findings suggest that continuously monitored pulmonary haemodynamics may inform clinical decision making, improve clinical outcomes and reduce healthcare costs in patients with heart failure.^16,17^ The annual per patient therapeutic cost for a patient with PAH is £20K/year rising to £100K/year for those on triple combination therapy.^18^ However, recent phase III studies demonstrate that discontinuation of regulatory approved therapies is not uncommon (21.7-29.1%^6,19^). Therefore, the ability to measure pulmonary artery pressures via an implantable pressure monitor may provide a means of matching individual patients with PAH to therapies effective for their disease and track changes in pressure with time to provide early, meaningful indicators that may be used to alter drug therapy prior to clinical deterioration.^20^ This approach may also provide an alternative means of undertaking phase II clinical studies in patients with PH.

The present study examines the safety, short and long-term accuracy of a novel pulmonary artery pressure monitor system, to detect potentially clinically relevant alterations in pulmonary artery pressures in animal models with raised pulmonary artery pressures.

## Methods

### Anaesthetic and vascular access

All animal studies were conducted in Yorkshire white pigs (25-43 kg) in accordance with The Animals (Scientific Procedures) Act 1986 under UK Home Office Project License 40/3695 and 40/3722. Under anaesthesia (intramuscular azaperone, 40 mg/mL at 6 mg/kg; intravenous propofol, 10 mg/mL at 3 mg/kg; and isoflurane, 2% to 3% in 100% O2 via endotracheal tube) the right femoral vein and artery (device implantation), or internal jugular and carotid artery (recatheterisation) were exposed using standard surgical techniques and vascular access gained via a 14F (Bard, USA) and 7F (Medtronic, UK) introducer respectively. With radiographic guidance (BV Pulsera, Philips, UK) pulmonary angiography and left and right heart catheterisation were performed using standard interventional techniques^21,22^ Invasive pressure was measured by fluid filled and Millar catheter (Ventri-Cath 510 and Mikro-Cath, Millar, USA) with simultaneous recording of ECG and core temperature using a PowerLab 8/35 (AD Instruments). Data were displayed and analysed using LabChart Pro (AD Instruments).

### Pressure monitor implantation

Prototype pulmonary artery pressure monitors and delivery systems were provided by Endotronix (Chicago, USA). The pulmonary artery pressure monitors were sanitised with 100% ethanol dip immediately prior to delivery system attachment using aseptic techniques. A 7F Swan-Ganz catheter (Edwards) was positioned in the left pulmonary artery via the right femoral vein. A 260cm 0.025’ Amplatz guidewire (Cook, UK) was inserted through the distal lumen and the Swan-Ganz removed. The pulmonary artery pressure monitors, attached to the distal end of a Delivery Catheter supported by a concentric Stability Sheath, were advanced to the target location over the guidewire (Figure 1). Pulmonary artery pressure monitors were positioned in the pulmonary artery or interlobar artery, orientated anterior-posterior and deployed by withdrawal of release wires attaching the Sensor to the Delivery Catheter (Figure 1). The Delivery Catheter and the guidewire were removed and a Millar catheter advanced through the Stability Sheath positioned proximal to the pulmonary artery pressure monitor for device calibration. Following implantation aspirin (37.5 mg) and clopidogrel (37.5 mg) were administered for 30 days.

**Figure 1.**
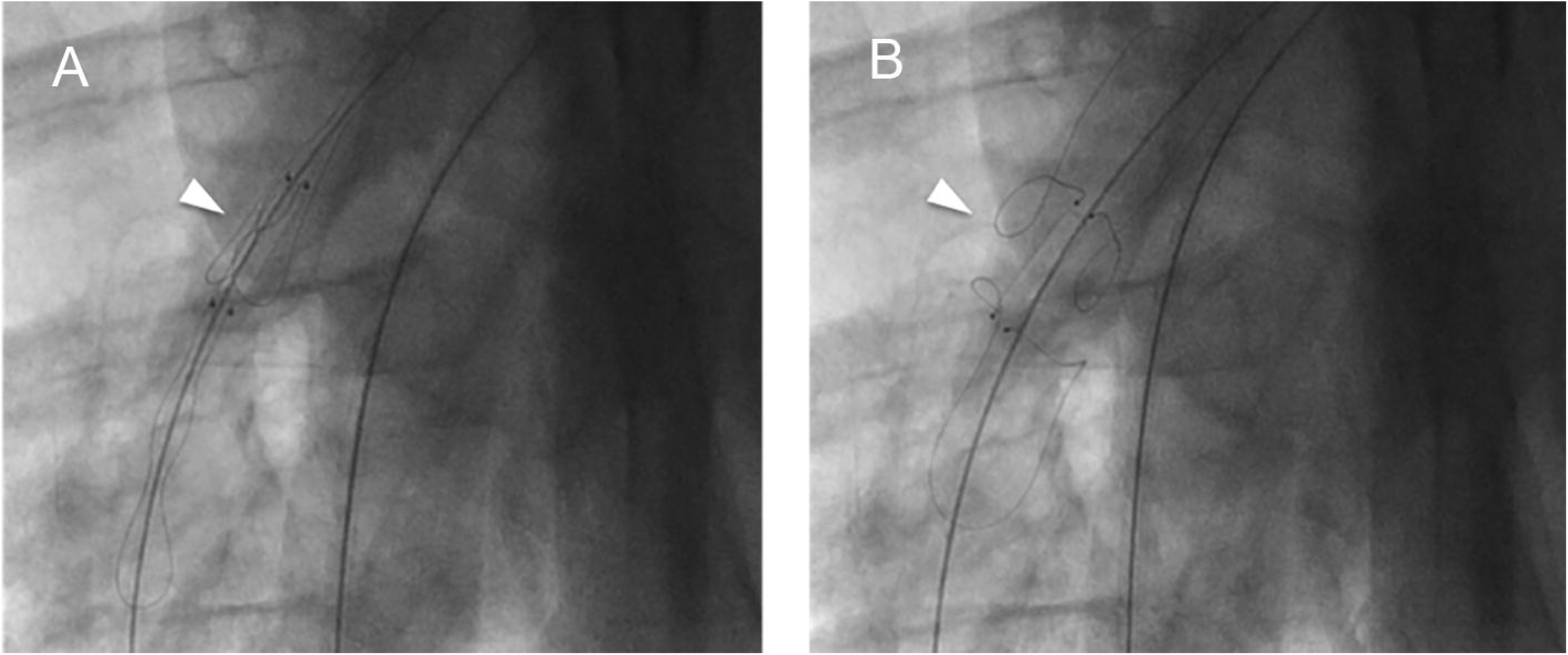
Representative fluoroscopic images of implantation. A) Implant at the target delivery location. The radiopaque markers on the implant body shows implant orientation within the vessel. B) Implant post-deployment with engaged anchors.

### Pressure monitor measurements

To ensure resting haemodynamic load, all readings were taken with animals anaesthetised (at implantation and right heart catheterisation) or sedated (for readings at time-points between implantation and repeat right heart catheterisation) with intramuscular injection of azaperone (40 mg/mL at 6 mg/kg). The reader was placed over the pulmonary artery pressure monitor, signal strength analysed and pulmonary artery pressure measured for 15 seconds.

### Echocardiography

Echocardiography was performed at baseline and following the induction of pulmonary hypertension (PH) (6S probe and Logiq E console, GE Healthcare). Images were acquired in the supine position via a parasternal window.

### Acute pulmonary hypertension

Following pulmonary artery pressure monitor implantation and calibration, thromboxane A2 agonist (TxA2, D0400, Sigma-Aldrich) was infused via the right femoral vein. The dose of infused TxA2 was increased at 5-minute intervals as previously described.^21-23^ Pulmonary artery pressure was measured using the implanted pulmonary artery pressure monitor (Endotronix, USA), fluid filled catheter and Millar catheter.

### Chronic pulmonary hypertension

Preliminary studies were undertaken to determine the optimal route and dose of monocrotaline administration (data not shown). Pulmonary artery pressure monitors were implanted 30 days prior to the induction of PH. At day 0 right heart catheterisation was repeated during which pressure monitors were calibrated to fluid filled measurements and PH induced by intraperitoneal administration of monocrotaline 20 mg/kg (Sigma-Aldrich, UK) at day 0 and haemodynamics assessed by repeat right heart catheterisation at day 30 (60 days after the implantation of the pulmonary artery pressure monitor).

### Histology

To demonstrate the interaction of the pulmonary artery pressure monitor with the vessel wall, histological samples were obtained following final catheterisation. As previously described animals were sedated by intramuscular injection of azaperone (6–8 mg/kg) and euthanised by intravenous injection of phenobarbital (40 mg/kg) in accordance with The Animals (Scientific Procedures) Act 1986 under UK Home Office Project License 40/3722.^21^ The chest was opened via a midline thoracotomy and the pleura and pericardium dissected to expose the heart and great vessels. The inferior vena cava, superior vena cava, and descending aorta were cross-clamped. Large bore, ridged cannula were placed in the pulmonary artery via an incision in the right ventricle and the left atrium via an incision in the left atrial appendage and secured with umbilical tape. The lungs were flushed with 3 L of 0.9% NaCl and perfusion fixed in 10% formalin. The heart and lungs were removed en-bloc, the airways filled with formalin and the sections suspended in 10% formalin for two weeks. Histological sections were dissected anatomically and preserved in 70% ethanol. Tissue sections were cut post-fixation, dehydrated in graded alcohols, and paraffin embedded. Five micrometer sections were mounted and stained with hematoxylin and eosin (H&E) and Verhoeff-van Gieson (VVG) to demonstrate the pulmonary artery pressure Sensor and anchors *in situ*. Images were acquired on an inverted light microscope (Olympus BX41) with a Leica camera analysis performed using Leica.

### Micro CT and Faxitron

After tissue fixation, microCT of pressure monitor and surrounding vessel was performed and data 3-D reconstructed (Skyscan 1172, Bruker).

### Statistical analysis

Data are expressed as mean ± S.E.M. with normality of distribution determined by Kolmogorov-Smirnov test and differences between data sets assessed using paired or unpaired student t-test or Mann-Whitney (unpaired) or Wilcoxon (paired) as appropriate in Prism 6.0 for Macintosh (GraphPad Software).

## Results

### Safety

No implant related adverse events were identified. Specifically, there was no evidence of perforation or dissection and no thrombus formation was identified angiographically or histologically.

### Implantable sensor provides accurate, acute measurement of pulmonary artery pressure

To determine the accuracy of the implanted pulmonary artery pressure monitor-derived, and reference catheter, pressure readings were made during the induction of acute PH in two Yorkshire White Swine. Real-time pulmonary artery pressure readings made by implanted pulmonary artery pressure monitor and gold standard Millar catheter were well matched (Figure 2A). Over a physiological range, pressure measured by implanted pulmonary artery pressure monitor and reference catheter was concordant with a bias of −0.43 (2.5 S.D., 95% CI −5.3 - 4.4, Figure 2B) demonstrating accurate acute measurement of pulmonary artery pressure.

**Figure 2.**
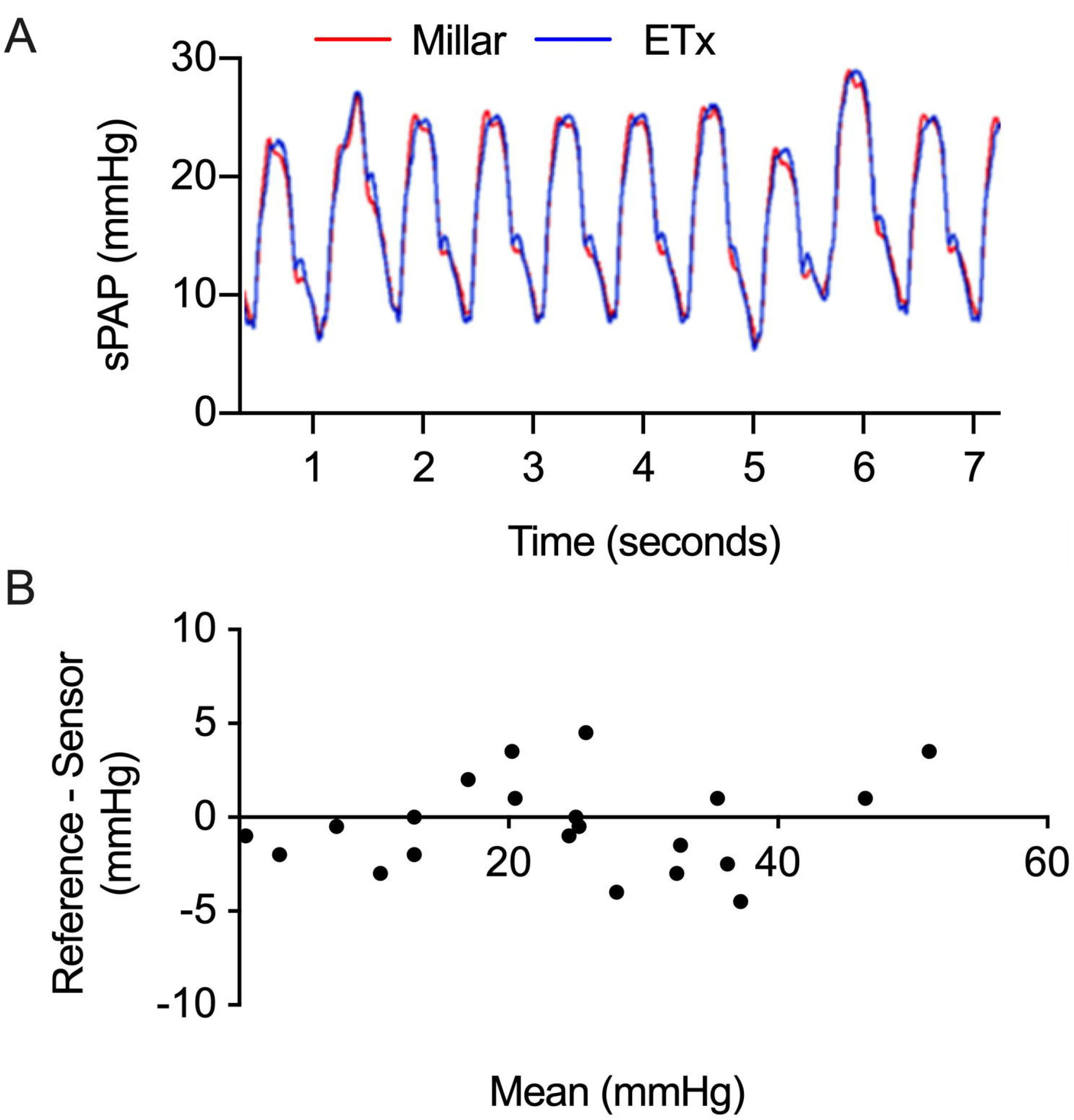
A) Pressure waveforms measured with the reference invasive pressure measurement (Millar) and the Sensor. B) Bland-Altman plot showing Millar and ETX pressure measurements.

### Intraperitoneal administration of monocrotaline results in chronic pulmonary hypertension and right heart failure

To determine long-term vascular compatibility and accuracy of pressure readings PH was induced in six animals by i.p. administration of monocrotaline MCT (20mg/kg). Five control animals were administered 0.9% normal saline solution. Administration of MCT led to increased dyspnoea, lethargy (subjective), and the development of dilated ear and abdominal veins. Consistent with increased pulmonary artery pressures, pulmonary artery acceleration time (PAAT) was decreased at day 30 (Figure 3A). This finding was confirmed by right heart catheterisation at day 30 demonstrated increased systolic pulmonary artery pressure in MCT treated animals compared to saline treated animals (23.1 +/- 1.0 vrs 31.74 +/- 1.4, p < 0.01) and compared to day 0 measurements in MCT treated animals (20.7 +/- 0.4 vrs 31.74 +/- 1.4, p < 0.01, Figure 3B). Furthermore, histological examination of the lungs demonstrated thickened, remodelled small pulmonary arterioles and examination of the liver demonstrated zone 2 and 3 acinus congestion with hepatocellular dropout, in keeping with the development of PH and right ventricular failure in MCT treated animals (Figure 3C).

**Figure 3.**
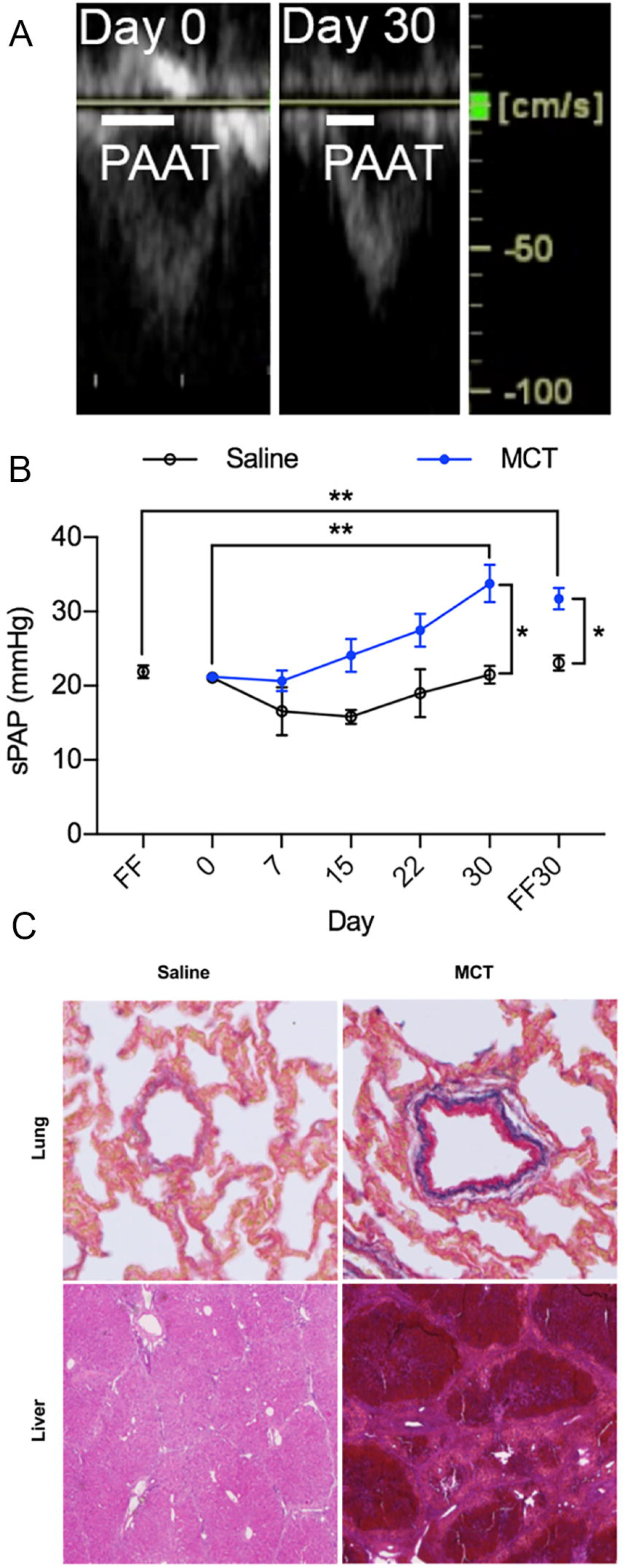
A) Representative Doppler echocardiography images of pulmonary artery flow showing pulmonary artery acceleration time (PAAT) showing decreased PAAT with the development of PH. B) sPAP measurements using Sensor and fluid-filled catheter over the course of the study showing chronic increase in sPAP over time (mean +/- S.E.M, *p < 0.05, ** p < 0.01, Mann-Whitney test or or Wilcoxon test as appropriate). C) Representative histopathological images of lung and liver sections from control and MCT treated animals. Administration of MCT resulted in increased small vessel muscularisation of the pulmonary arteries (Lung - Alcian blue elastic Van Gieson, 20x magnification) and zone 2 and 3 acinus congestion with hepatocelular dropout, in keeping with the development of PH and right ventricular failure (Liver – haematoxylin and eosin, 2.5x magnification).

### Implantable sensor stability and long-term pulmonary artery pressure measurement

Pulmonary artery pressure monitor position remained stable throughout the study with no identifiable longitudinal migration or rotation and no structural defect of the device (Figure 4). which was fully endothelialised (Figure 5). The increased pulmonary artery pressures measured by fluid filled catheter matched those measured by implanted pulmonary artery pressure monitor at day 0 and day 30 (Figure 3B). Pulmonary artery pressure monitor derived readings at day 15 to day 30 demonstrated a progressive increase in systolic pulmonary artery pressure (Figure 3B).

**Figure 4.**
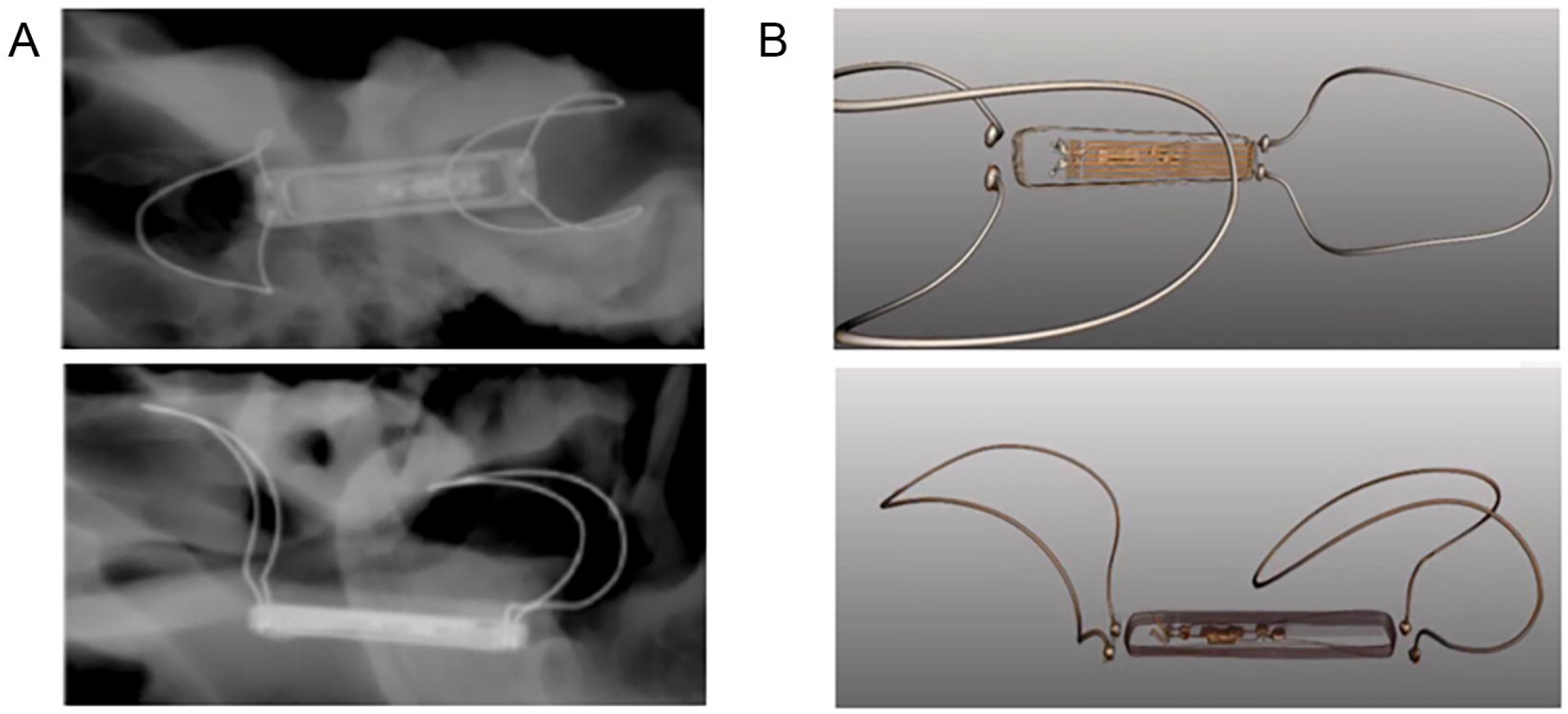
A) Micro-CT images of the prototype implant from different views. B) Reconstructed 3D computer model based on the micro-CT images.

**Figure 5.**
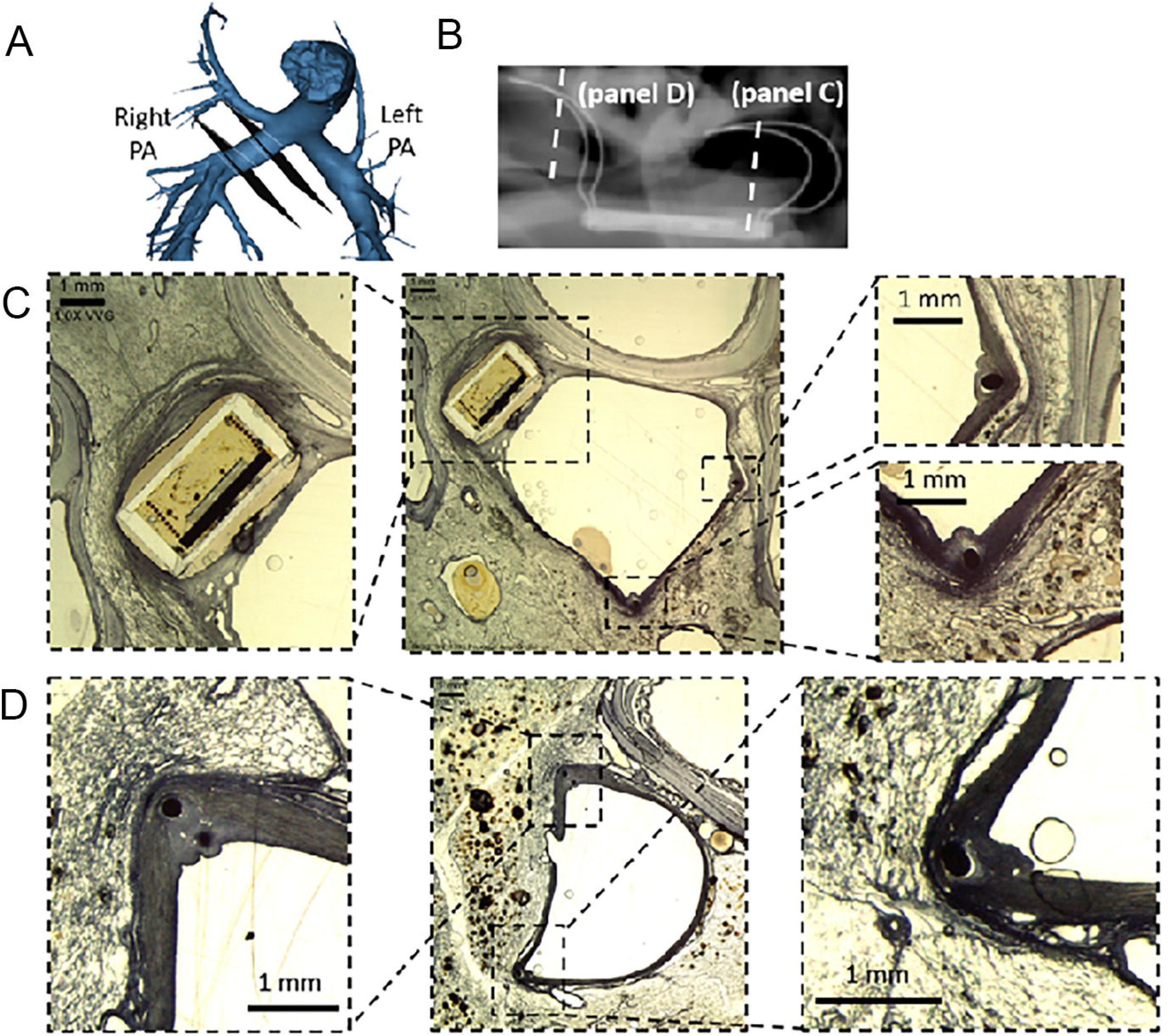
Histology. A and B – location of sensor and histological section within the pulmonary artery. Hematoxylin and eosin (C) and Verhoeff-van Gieson (D) stained sections of sensor body and anchors demonstrating endotheliasaiton.

## Discussion

The present study demonstrates the pre-clinical safety, vascular compatibility and accuracy of a novel telemetry based pulmonary artery pressure monitor system in acute and chronic large animal models of PH. Real-time pressure wave form traces were well matched to gold-standard reference catheter measurements, and device-measured pressure readings tracked the development of disease in chronic models over a 30-day period.

Right heart catheterisation is an essential diagnostic investigation of patients with suspected pulmonary hypertension and may be used in selected patients to assess the response to treatment or guide the requirement for advanced therapies or transplantation. Beyond a diagnostic role, haemodynamic parameters aid risk stratification of patients with PAH and guide choice of therapy.^2,4,24^ Therapeutic options for patients with PAH and CTEPH have expanded over recent years, however beyond those with an acute vasodilator response who have excellent survival with calcium channel antagonists,^25^ there is little to guide physicians to which drugs may be best suited to individual patients and limited evidence based means by which to determine response to therapy.^2^ Options for patients with other forms of PH are limited.

PAH specific therapies alter pulmonary haemodynamics including mean pulmonary artery pressure. Pooled patient-level analysis of clinical trial data have demonstrated that changes in pulmonary haemodynamics, with knowledge of treatment allocation, is predictive of clinical events.^10^ The development of implantable pulmonary artery pressure monitors offers the potential to determine pulmonary artery pressure with greater frequency than previously possible without the need for invasive catheterisation. In patients with heart failure due to left heart disease the benefit of haemodynamic parameter-guided therapy has been studied and elevation of cardiac filling pressures has been shown to precede clinical events by ≥1 to 2 weeks.^11,12^ More recently, early identifiable physiological changes have been detected using implantable pulmonary artery pressure monitors and proactive pressure-guided heart failure management has been demonstrated to reduce hospitalisation in comparison to standard care in patients with heart failure and those with heart failure and coexisting lung disease.^13-15^ As such it is possible that pulmonary artery pressure monitoring and proactive pressure-guided heart failure management may provide benefits to groups of patients with PH without currently approved therapies. In patients with all forms of PH, changes in haemodynamics may be driven by a range of factors such as pulmonary vascular remodeling and constriction, fluid status, hypoxia, infection, therapeutic adherence, drug interactions or physical activity. Remote monitoring of PA pressure and additional parameters used in routine clinical practice has the potential to facilitate differentiation of cause of decompensation, aiding instigation of appropriate, timely management.^26^

Furthermore, there is limited long-term data on variability of pulmonary artery pressure in patients with PAH.^27^ Factors such as patient position and haemodynamic loading,^28–30^ hypoxia^31^ and exercise^32,33^ have been shown to alter pulmonary artery pressure in short-term studies. The development of pulmonary artery pressure monitors that provide long-term haemodynamic data provides an opportunity to investigate physiological and clinical questions essential to the understanding of disease and provides the opportunity to assess therapeutic interventions in clinical studies of novel agents.

